# Structural basis of the differential binding of engineered knottins to integrins αVβ3 and α5β1

**DOI:** 10.1101/565358

**Authors:** Johannes F. Van Agthoven, Hengameh Shams, Frank V. Cochran, José L. Alonso, James R. Kintzing, Kiavash Garakani, Brian D. Adair, Jian-Ping Xiong, Mohammad R. K. Mofrad, Jennifer R. Cochran, M. Amin Arnaout

## Abstract

Targeting both integrins αVβ3 and α5β1 simultaneously appears to be more effective in cancer therapy than targeting each one alone. The structural requirements for bispecific binding of ligand to integrins has not been fully elucidated. RGD-containing knottin 2.5F binds selectively to αVβ3 and α5β1, whereas knottin 2.5D is αVβ3-specific. To elucidate the structural basis of this selectivity, we determined the structures of 2.5F and 2.5D as apo-proteins and in complex with αVβ3, and compared their interactions with integrins using molecular dynamics simulations. These studies show that 2.5D engages αVβ3 by an induced fit, but conformational selection of a flexible RGD loop accounts for high affinity selective binding of 2.5F to both integrins. The contrasting binding of the highly flexible low affinity linear RGD peptides to multiple integrins, suggests that a “Goldilocks zone” of conformational flexibility of the RGD loop in 2.5F underlies its selective binding promiscuity to integrins.

## Introduction

Heterodimeric α/β integrins comprise a large family of divalent cation-dependent adhesion receptors that mediate cell-cell and cell-matrix interactions, which underlie their essential roles in normal metazoan physiology but also in contributing to many diseases including pathologic thrombosis, inflammation, autoimmunity and cancer (Raab-Westphal et al., 2017). In response to cell-activating stimuli, intracellular signals are generated that rapidly convert integrins into a ligand-competent state, a process termed inside-out signaling (Arnaout et al., 2007). Physiologic ligands, prototyped by the Arg-Gly-Asp (RGD) sequence motif, then bind the integrin head (formed of the α-subunit propeller and β-subunit βA domains) (Xiong et al., 2002). A carboxylate group of the ligand Asp makes an electrostatic contact with a Mg^2+^ ion coordinated at a *m*etal-*i*on-*d*ependent *a*dhesion *s*ite (MIDAS) of the βA domain, and the ligand Arg inserts into a pocket in the α-subunit propeller. Ligand binding induces tertiary changes in βA that are converted to quaternary changes in the ectodomain, thus forging links of the integrin cytoplasmic tails with the actin cytoskeleton to regulate cell function, a process termed outside-in signaling (Friedland et al., 2009).

Therapeutic targeting of integrins has generally focused on development of peptides or small molecules that primarily target a single integrin (Kapp et al., 2017), an approach that has been effective in platelets, where integrin αIIbβ3 is most abundant (Coller and Shattil, 2008). However, in other cell types expressing multiple integrins, high selectivity for a single integrin may promote upregulation of another integrin sharing the same ligand, leading to reduced effectiveness, drug resistance, or even paradoxical effects. This scenario may be particularly relevant in cancer cells, where primary targeting of αVβ3 with cilengitide failed to prolong survival of patients with glioblastoma (Mason, 2015), likely related to unfavorable pharmacokinetics, enhanced α5β1-mediated cell migration (Caswell et al., 2008; Christoforides et al., 2012) and agonist-like behavior (Reynolds et al., 2009). Recent studies also showed superiority of targeting αVβ3 plus α5β1 as compared to αVβ3 alone in noninvasive *in vivo* imaging of brain cancer in mice (Moore et al., 2013).

The engineered 3.5kDa miniproteins knottins 2.5D and 2.5F bind with nanomolar affinity to αVβ3 (2.5D) or to both αVβ3 and α5β1 (2.5F) (Kimura et al., 2009a). 2.5D and 2.5F only differ in four residues: two on either side of the RGD motif (Figure 1A). In this report, we determined the solution structures of 2.5F and 2.5D and their crystal structures in complex with αVβ3. Our results show that the 2.5F and 2.5D use different binding modes to interact with αVβ3 that are critically dependent on the degree of conformational flexibility of the respective RGD loop backbone. These data suggest that flexibility of the RGD loop in 2.5F is just sufficient to allow it to bind both integrins by adopting conformations to fit both binding sites but not so large, as in linear RGD peptides, that the entropic cost of stabilizing the loop in a single conformation will compromise its high-affinity binding.

**Figure 1.**
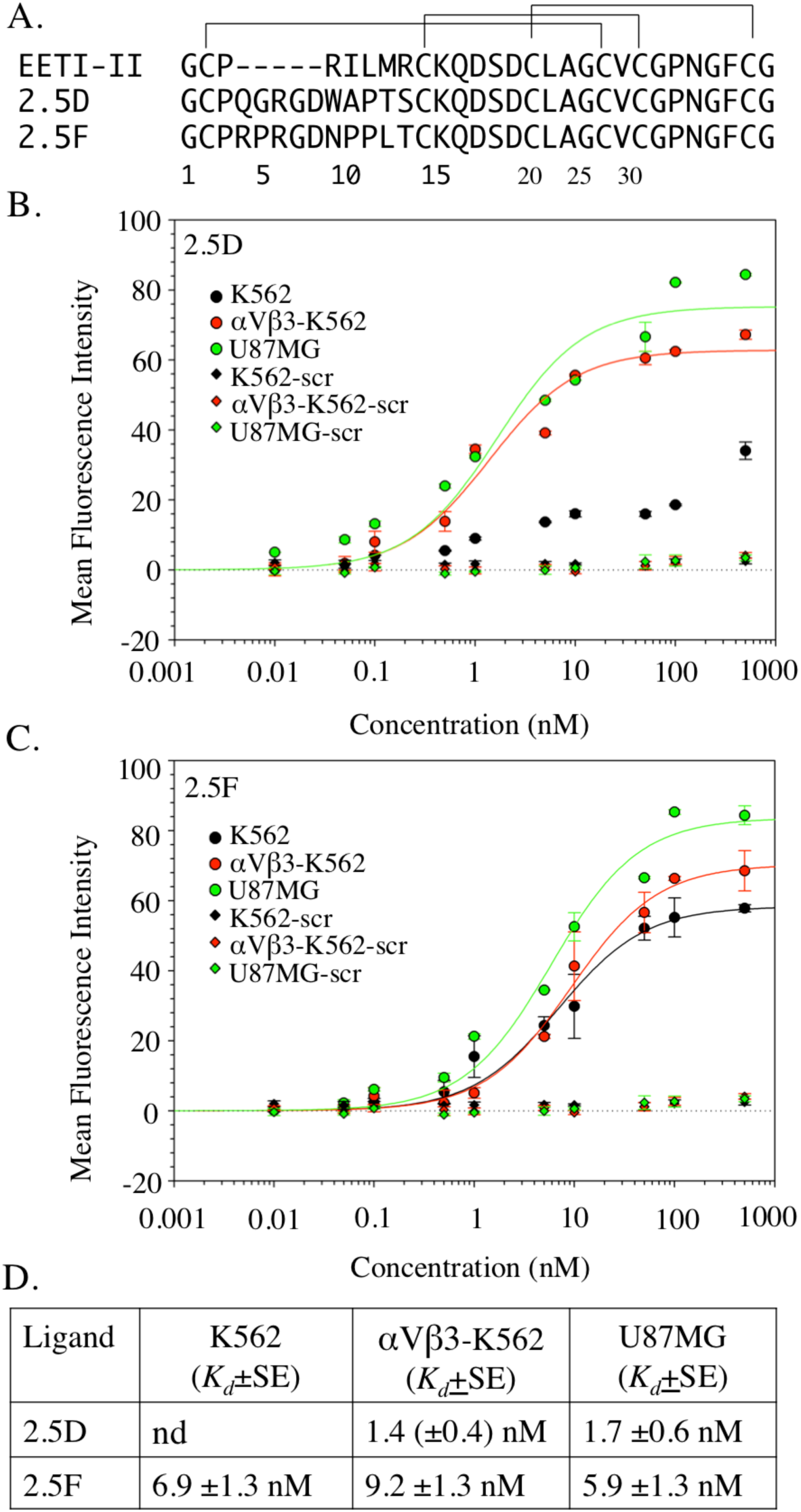
Primary sequence and binding properties of knottins 2.5D and 2.5F. (A) The primary structure of EETI-II and of the engineered knottins 2.5D and 2.5F, where the 6-residue trypsin-binding sequence (P^3^RILMR) in native knottin is replaced with an 11-residue sequence containing the RGD motif. (B, C) Dose-response curves showing binding of Fc fusions of 2.5D (B), 2.5F (C) or a scrambled (scr) knottin (B, C) to native integrins expressed on U87MG, αVβ3 expressed on transfected K562 cells (αVβ3-K562), and native α5β1 on K562 cells. Points display the mean and standard deviation for triplicate determinations. (D) Equilibrium binding constant (*K*_*d*_) values from the binding data shown in B, C, along with the standard error derived from the fitted curves. The *K*_*d*_ value derived from the curve fit of 2.5D-Fc binding to K562 in Figure 1B is not reliable as determined by the P value for the parameter, and thus not reported in Fig.1D. nd, not determined.

## Results

### Integrin binding to 2.5D and 2.5F

We measured binding of 2.5D-Fc or 2.5F-Fc fusion proteins to K562 cells stably expressing recombinant αVβ3 (αVβ3-K562) and to K562 cells, which constitutively express α_5_β_1_ integrin. 2.5D bound to αVβ3-K562 cells with nanomolar affinity (1.4 ± 0.4 nM, Figure 1B, D), but exhibited no measurable binding affinity to K562 cells (Figure 1B, D), over the same range, as was also true for the scrambled knottin FN-RDG2 (where the RGD motif is replaced with RDG (Kimura et al., 2009b)). In contrast, 2.5F bound both αVβ3-K562 and K562 with similar affinities (6.9 ± 1.3nM and 9.2 ± 1.4 nM, respectively) (Figure 1C, D. Both 2.5D and 2.5F also bound U87MG glioblastoma cells (1.7 ± 0.6 nM and 5.9 ± 1.3 nM, respectively) (Figure 1B-D), which express high levels of αVβ3 (Dumont et al., 2009).

### NMR structures of 2.5D and 2.5F

To begin to elucidate the structural basis for selectivity of knottin/integrin binding, we first determined the solution structures of 2.5D and 2.5F by NMR (Table 1). As expected, both knottins assumed the same compact structure held together by three disulfide bonds, typical of the cysteine inhibitor family (Figure 2A, B). However, structure of the engineered RGD-containing loop flanked by prolines 3 and 11 was drastically different in the two knottins (Figure 2A, B). In 2.5D, this loop maintains a nearly single packed conformation (Figure 2A, C, D), but is flexible in 2.5F (Figure 2B-D).

**Table 1.**
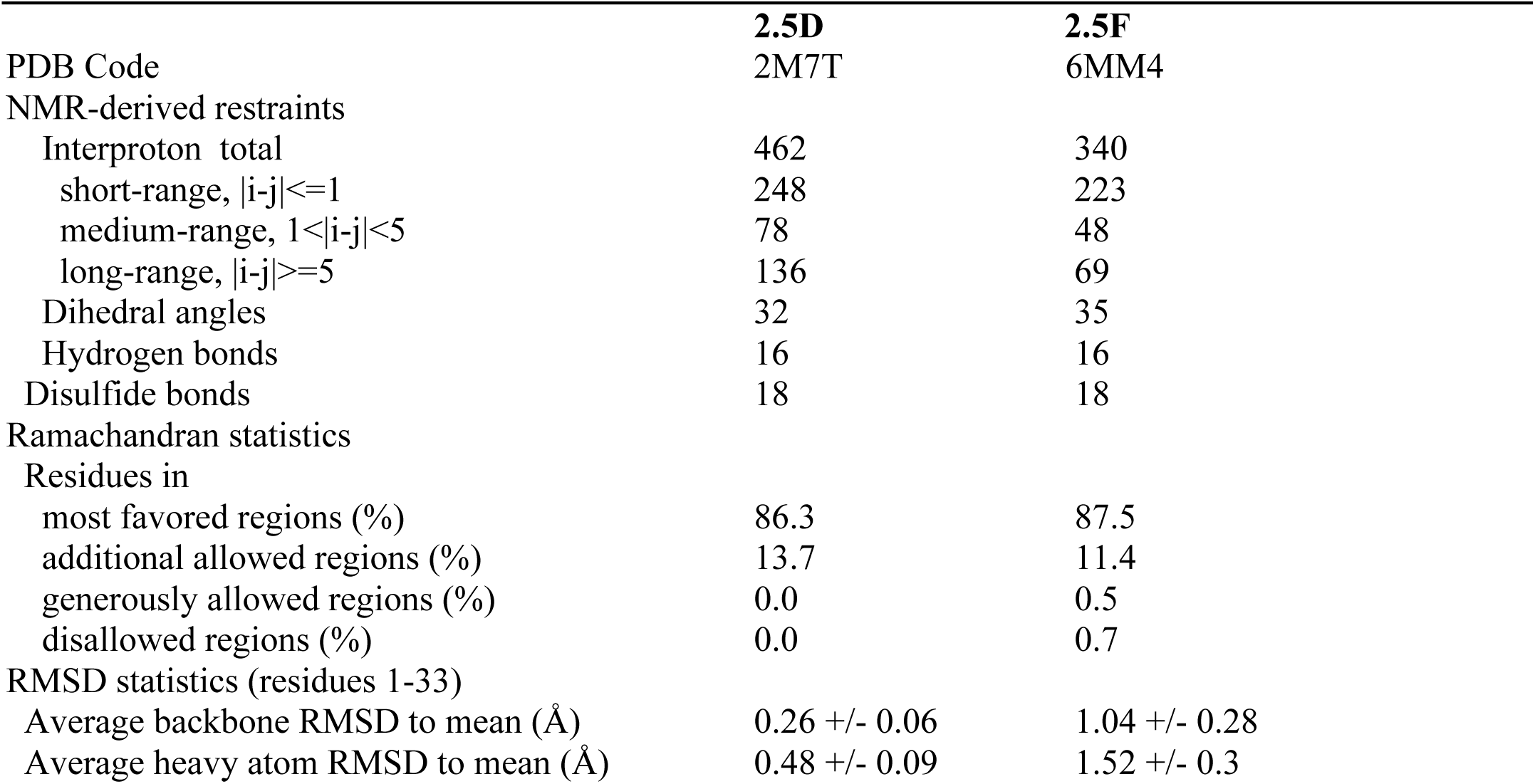
NMR structural statistics for 2.5D and 2.5F 20-conformer ensembles.

**Figure 2.**
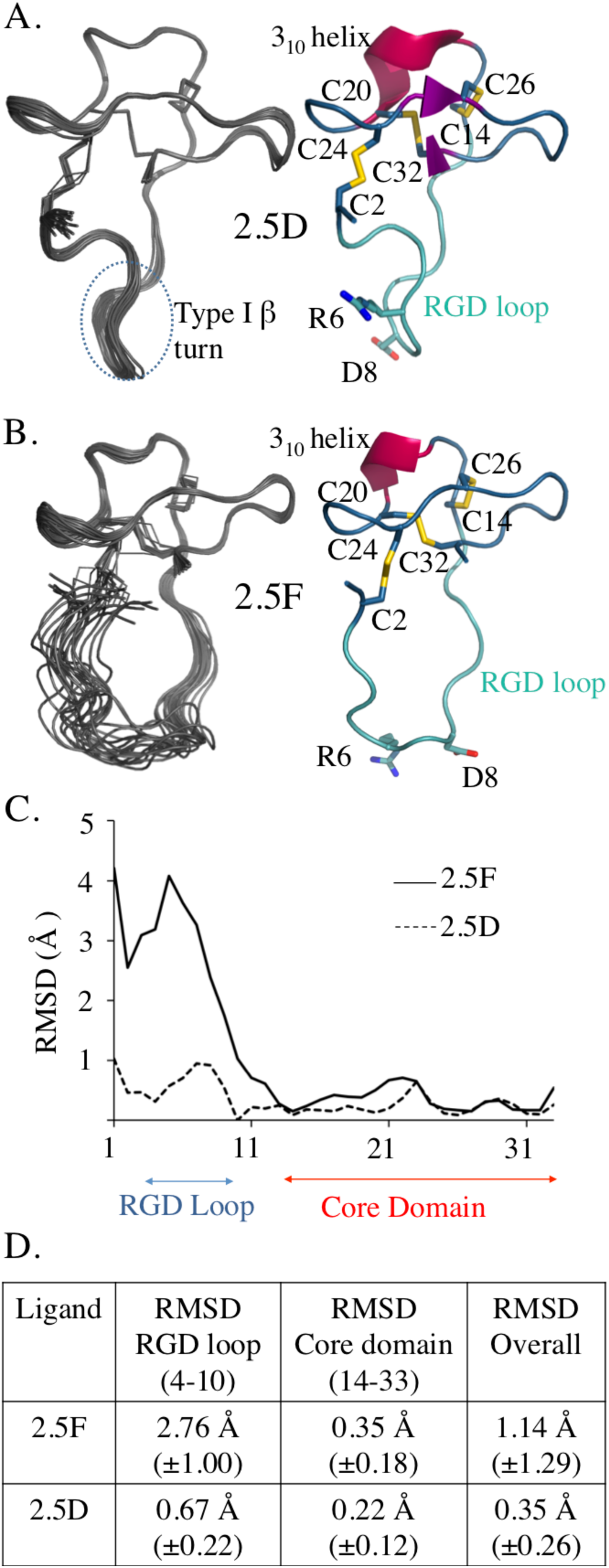
NMR structures and analysis of 2.5D and 2.5F. (A, B) Solution NMR structures of knottins 2.5D (A) and 2.5F (B) represented as the 20 lowest energy conformers. Ribbon diagrams of the lowest energy conformer of 2.5D and 2.5F are shown to the right. The RGD loop is in cyan with the integrin RGD binding sequence in sticks. The core domain is blue with disulfide bonds in yellow sticks. The 3_10_ helix is represented in red cartoon. (C) RMSD of solution structure of 2.5D and 2.5F plotted versus residue number. The RGD loop (residue 4-10) and core domain (residue 14-33) are indicated. (D) Table summarizing the RMSD values per domain shown as mean±S.D.: core domain (residue 14-33) and RGD loop (residue 4-10).

### X-ray structures of integrin-bound 2.5D and 2.5F

Knottins 2.5D and 2.5F were each soaked into preformed αVβ3 ectodomain crystals in presence of 1 mM Mn^2+^ and the crystal structure of the respective complex was determined as previously described (Van Agthoven et al., 2014; Xiong et al., 2002). Simulated annealing composite omit maps showed clear ligand density, and allowed complete tracing of the knottin macromolecule (Figure 3, Table 2, Figure S1) with Real Space Cross Correlations (RSCCs) for RGD in both knottins of 0.93-0.97, suggesting almost full occupancy of the ligand. The RGD motif of each ligand inserts into the crevice between the propeller and βA domains and contacts both in an identical manner (Figure 3A-F). The Arg^6^ guanidinium of each knottin contacts α_V_-Asp^218^ of the propeller, with a carboxylate from Asp^8^ contacting the MIDAS Mn^2+^. In the αVβ3/2.5D structure, 2.5D residues Ala^10^, Pro^11^, Pro^28^, Asn^29^ and Phe^31^ form additional van der Waals contacts with βA (Figure 3G). In the αVβ3/2.5F structure, 2.5F-Arg^4^ hydrogen bonds βA-Asn^313^ and contacts the ADMIDAS metal ion indirectly through a chloride ion (Figure 3F). Other interactions include van der Waals contacts of 2.5F residue Pro^10^, Pro^11^, and Phe^31^ with βA (Figure 3H). These interactions, which bury a surface area of 654.6 Å^2^ for αVβ3/2.5F and 606.2 Å^2^ for αVβ3/2.5D structures account for the high affinity binding of each ligand to αVβ3.

**Figure 3.**
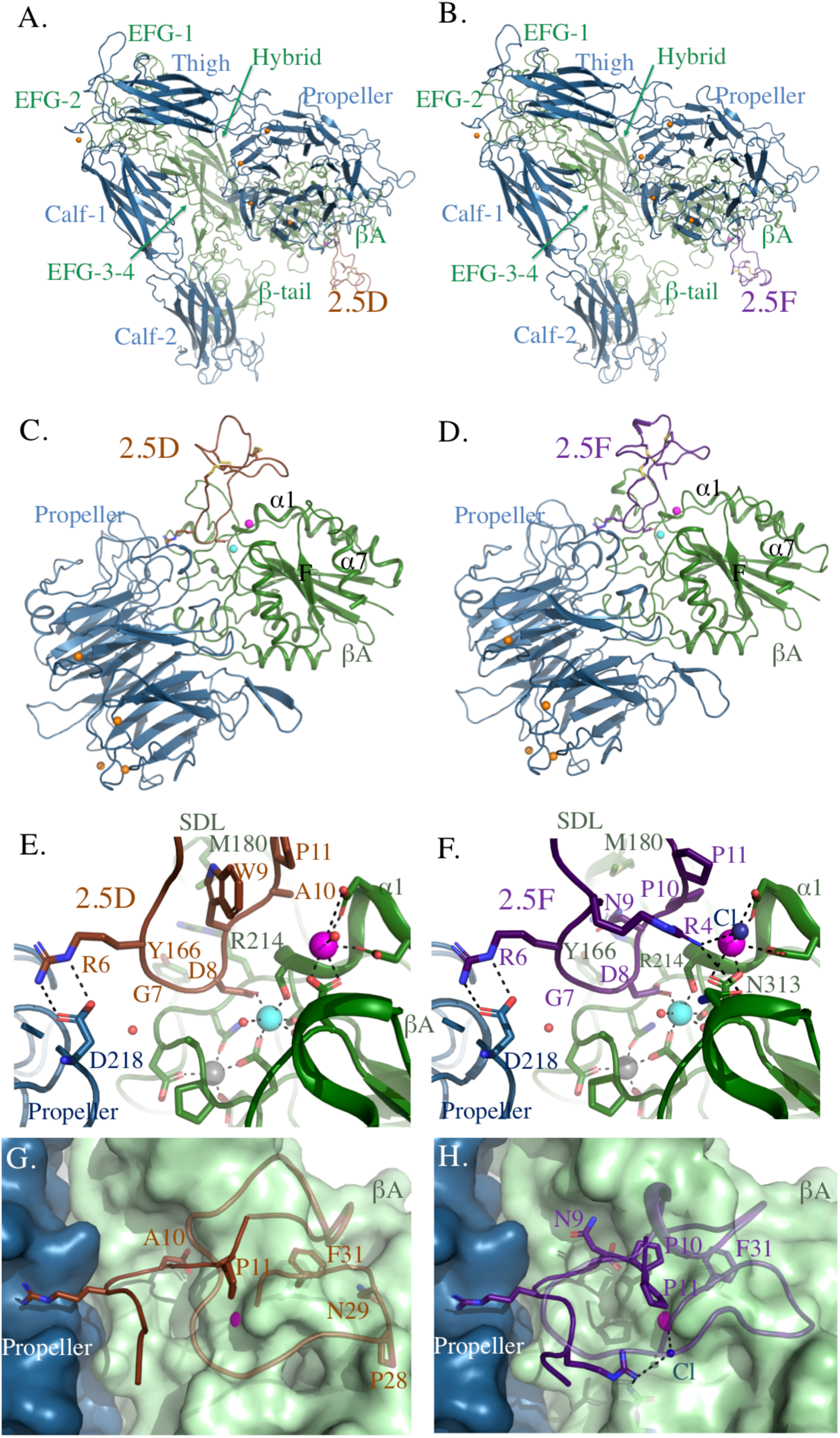
Crystal structures of αVβ3 bound to knottins 2.5D or 2.5F. (A, B) Ribbon diagrams of αVβ3 (αV is in blue and β3 in green) bound to 2.5D (brown in A, C, E and G) or 2.5F (purple, in B, D, F and H). (C, D) Ribbon diagrams of the αVβ3 head bound to 2.5D (C) and 2.5F (D). The propeller is in blue and βA domain in green. Mn^2+^ ions at LIMBS (gray), MIDAS (cyan) and ADMIDAS (magenta) are shown as spheres (also in E-H). (E, F) Ribbon diagrams showing key electrostatic and hydrogen bond interactions and metal ion coordination in the structure of αVβ3/2.5D (E) and αVβ3/2.5F (F). 2.5F-R4 hydrogen bonds with β3-N313. Additionally, 2.5F contacts the ADMIDAS ion through a chloride (Cl) ion represented as a blue sphere. Water molecules are shown as small red spheres. (G, H) Solvent accessible surface view of the integrin/ligand interface showing residues in 2.5D (A10, P11, P28, N29, and F31) and in 2.5F (N9, P10, P11 and F31) forming van der Waals (≤4Å) contacts with the βA domain. See also Figure S1 and S2.

**Table 2.**
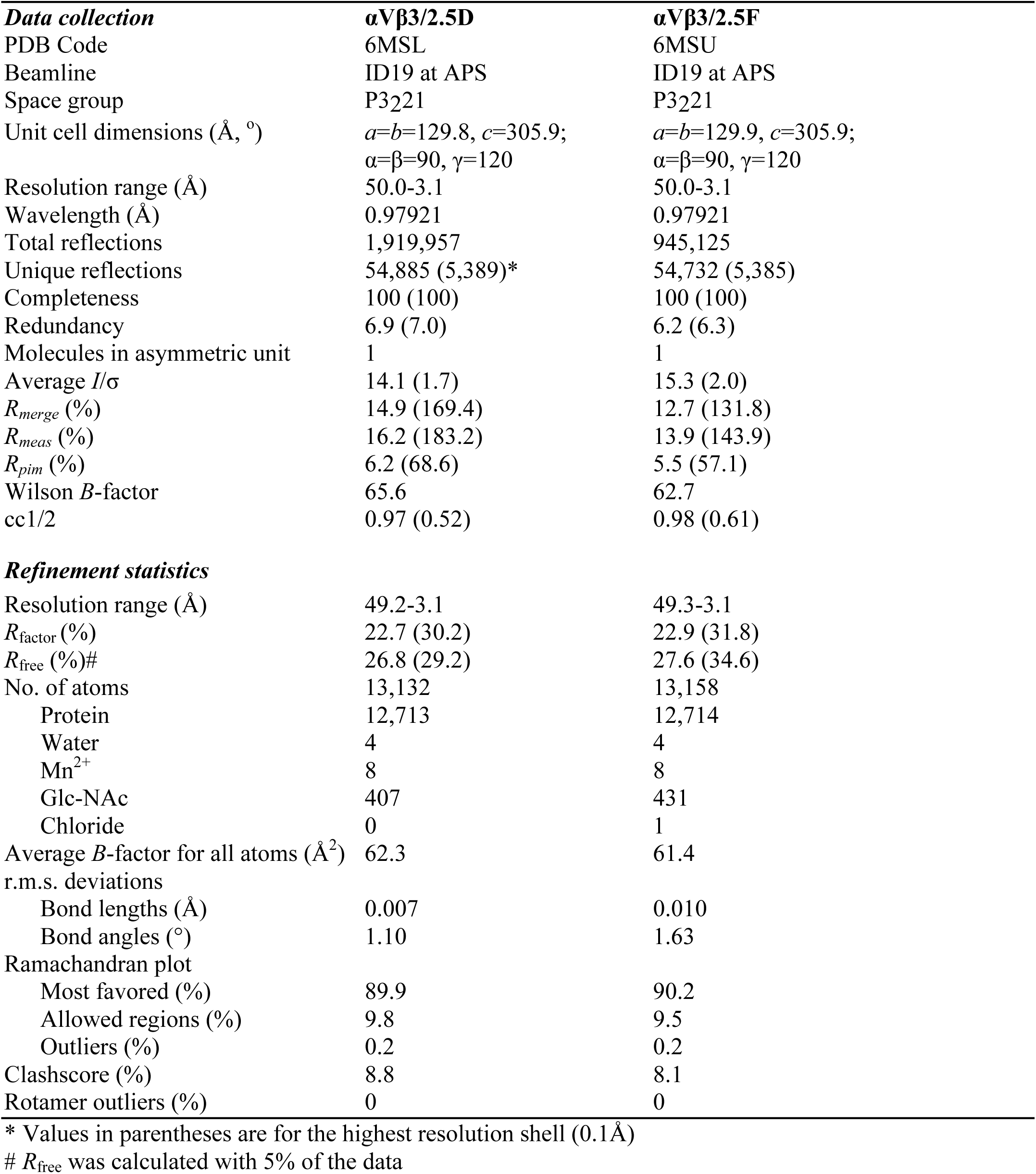
Data collection and refinement statistics.

As with binding of the natural ligand FN10 or the partial agonist cilengitide to αVβ3 (Van Agthoven et al., 2014; Xiong et al., 2002), binding of 2.5D or 2.5F induced a 3.7Å inward movement of the α1 helix of the βA domain towards the MIDAS Mn^2+^, and restructuring of the F/α7 loop (Figure S2A), confirming that both knottins are partial agonists. The shape of the CD loop of the βTD in both knottin/integrin structures was also comparable to the one published for αVβ3/wtFN10 (Van Agthoven et al., 2014) (Figure S2B-D). However, whereas the crystal structure of the pure antagonist hFN10 bound to αVβ3 showed a h-bond between β_3_-Glu^319^ of βA and β3-Ser^674^ of βTD and a visible glycan at Asn^711^, both features were absent in the αVβ3/2.5F and αVβ3/2.5D structures (Figure S2B-D).

### Conformations of the RGD-containing loops of 2.5D and 2.5F bound to αVβ3

In contrast to the major differences in conformation of the RGD-containing loops of 2.5D and 2.5F (Figure 2), the two loops were largely superposable in the integrin-bound state (Figure 4A), with a root mean square deviation (r.m.s.d.) of 0.62±0.27Å^2^ (mean±sd). Superposing the crystal structures of the integrin-bound loops on the respective NMR structure of the lowest energy state showed dramatic differences in the RGD-containing loop of 2.5D (Figure 4B). The r.m.s.d. of this loop in 2.5D between the integrin-bound and solution states is 3.52±1.01Å^2^, which is significantly higher than its narrow r.m.s.d. in solution that is maintained in all 20 conformers (0.67±0.22 Å^2^, Figure 2D, Figure S3). The apo-protein state is stabilized by a 4-residue type I β-turn spanning Arg^6^ to Trp^9^, with hydrogen bonds involving the carbonyl oxygens and amide nitrogens of Trp^9^ and Arg^6^, and maintains a distance of ∼6.4 Å between the β carbons of Arg^6^ and Asp^8^ 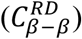(Figure 4C). The importance of a Trp residue immediately after RGD in forming a β-turn was previously noted (Park et al., 2002). When 2.5D is bound to αVβ3, the β-turn unfolds, with Trp^9^ moving from the solvent-exposed state to form an internal van der Waals bond with Gly^5^ (Figure 4D), thus extending the 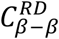 distance to 9Å (Figure 4E). In contrast, the r.m.s.d. of the RGD-containing loop of 2.5F between the bound and solution structure is 3.13±1.64Å^2^, comparable to its r.m.s.d. in solution (2.76±1.0 Å^2^) (Figure 4F), with conformational flexibility of the RGD loop backbone reflected in 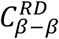 varying from 4.1Å in the lowest energy state to 8.8Å in conformer #14 (Figure 4G, Figure S3) that approaches the 9.1Å 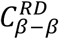 found in αVβ3/2.5F (Figure 4H) or RGD-bound α5β1 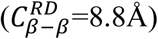 structures (Figure 4 I).

**Figure 4.**
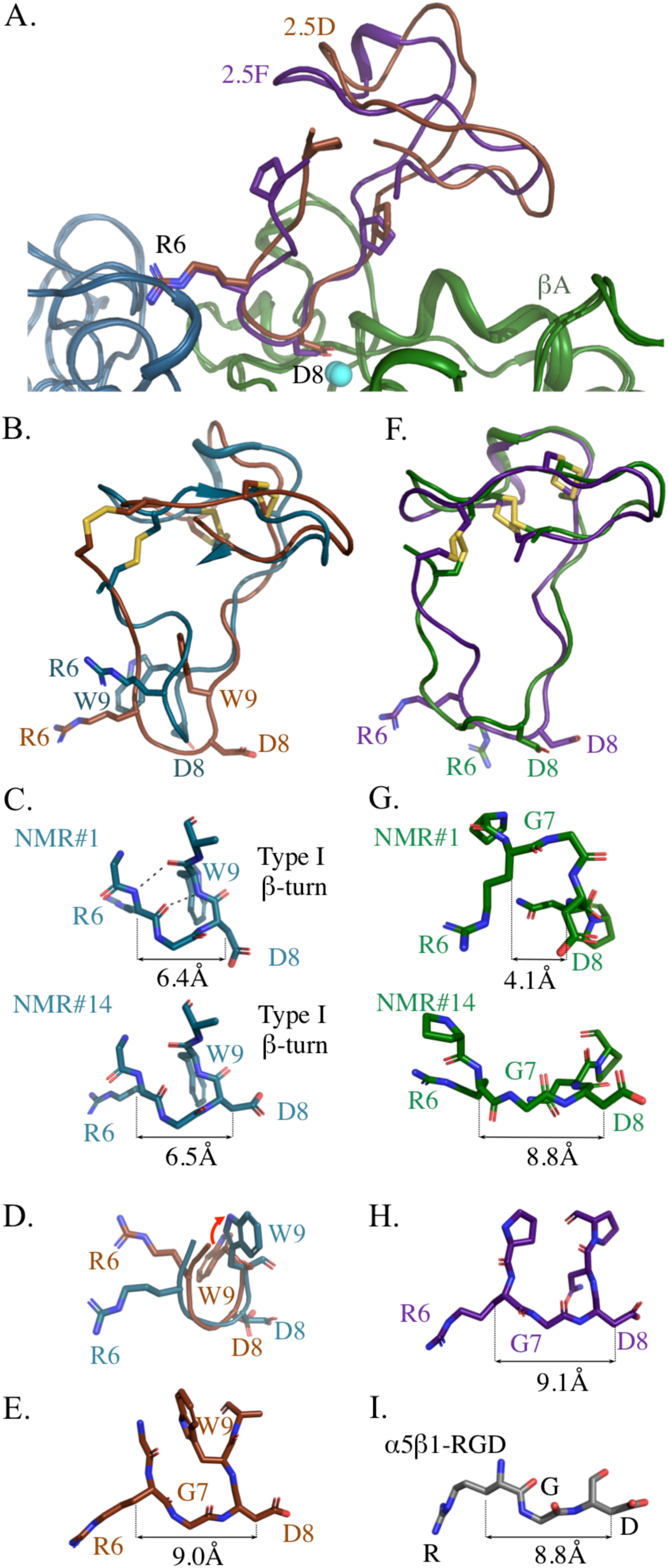
Structural comparisons of 2.5D and 2.5F in solution and integrin-bound. (A) Ribbon diagram of the superposed crystal structures of 2.5D (brown) and 2.5F (purple) in the integrin bound state. (B, F) Crystal structures of 2.5D (brown) and 2.5F (purple) superimposed on lowest energy model #1 of apo 2.5D (B, tale) and apo 2.5F (F, green). (C, G) Stick diagrams of the RGD loop in apo 2.5D (tale) and apo 2.5F (green) of lowest energy NMR #1 and of NMR#14. The 2.5D apo structure shows a β hairpin Type I turn in the RGD loop. Cβ-Cβ distances is stable among 2.5D conformers but is variable in case of 2.5F. (D) Superposition of GRGDW (residues 5-9) of 2.5D in the apo (tale) and bound states (brown). The red arrow shows conformational change of 2.5D-W9 from solvent-exposed state in the apo form to buried state in the crystal structure. (E, H) Crystal structures of the integrin-bound RGD loops of 2.5D (E) and 2.5F (H). (I) Stick diagram of the RGD loop bound to α5β1 crystal structure (PDB code 4WK4). The Cβ-Cβ distances between R6 and D8 in C, G, E, H, and R and D in I are shown. See also Figure S3.

### MD simulations of knottin binding to αVβ3 and α5β1

MD simulation was used to characterize the early stages in binding of 2.5F and 2.5D to αVβ3 and α5β1. The lowest energy NMR structures of 2.5F and 2.5D were docked onto the αVβ3 and α5β1 crystal structures, resulting in four protein complexes. In each complex, the knottin was moved 9Å away from the integrin surface allowing several water layers to form between the two before simulation was initiated. Over a 500-ns run, 2.5F associated with both αVβ3 (135±37 kcal/mole) and α5β1 (81±51 kcal/mole) (Figure 5A, B, Supplemental Movies 1 and 2), recapitulating the cell-based data (Figure 1 B-D). The first dual contact of 2.5F with αVβ3 was detected at 0.02 ns of simulation by 2.5F-Arg^6^ hydrogen-bonding β3-Thr^212^ and salt-bridging β3-Asp^150^, and 2.5F-Asp^8^ salt-bridging β3-Arg^214^ (Fig 5C). The first dual contact of 2.5F with α5β1 was detected at 0.35 ns, but the interaction stabilized at 1.25 ns through a salt-bridge between 2.5F-Asp^8^ and β1-Lys^182^ (Fig 5D).

**Figure 5.**
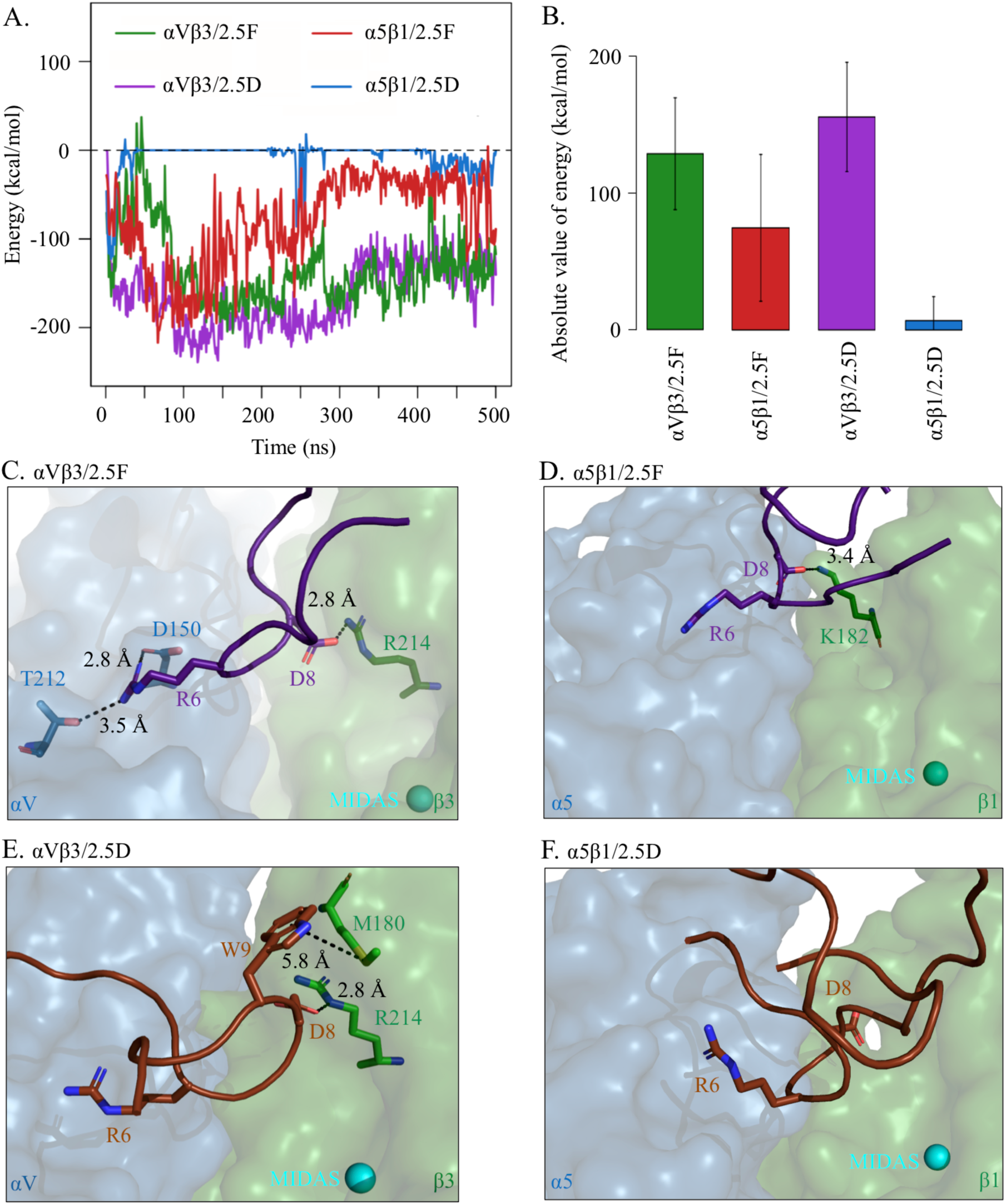
MD simulations of 2.5D/2.5F binding to αVβ3/α5β1. Knottins were initially separated by 9Å. (A) Energy over time and (B) bar graph (mean± SD) showing absolute value of binding energies between 2.5D and 2.5F to αVβ3 and α5β1 averaged over 500 ns of simulation time. Difference between 2.5D/α5β1 and 2.5D/αVβ3, 2.5F/αVβ3, 2.5F/α5β1 is significant at p<2.2×10^-16^. (C, D) Selected residues in structures of MD simulation of 2.5F binding to αVβ3 at t=0.020 ns (C) and to α5β1 at t=1.250 ns (D). (E, F) Selected residues in structures of MD simulation of 2.5D binding to αVβ3 at t=0.960 ns (E), and to α5β1 at t=1.250 ns (F). Distances (dotted lines) are indicated. The head segment of the respective integrin is displayed. See also Supplemental Movie 1, 2, 3 and 4.

As expected, 2.5D also bound αVβ3 effectively (164±37 kcal/mole) (Figure 5A, B, Supplemental Movie 3). The low energy of interaction between 2.5D-Trp^9^ and 2.5D-Gly^5^ suggested that surrounding residues in the binding pocket of αVβ3 are involved in the conformational switch of 2.5D-Trp^9^ from the solvent to the buried state. Consistently, over the course of the αVβ3/2.5D simulation, 2.5D-Trp^9^ first formed an S-π bond with β3-Met^180^ at 0.38 ns after the start of simulation, which was reinforced at 0.96 ns via an electrostatic interaction between 2.5D-Asp^8^ and β3-Arg^214^ (Figure 5E). In contrast 2.5D rapidly diffused away from α5β1 after 1.1 ns of interaction (7±19 kcal/mole) (Figure 5A, B), unable to sustain the initial binding of 2.5D-Arg^6^ to β1-Gln^221^ and β1-Asp^227^ at 0.01 ns (not shown) with additional contacts to the integrin (Figure 5F, Supplemental Movie 4).

### Binding of 2.5D and 2.5F to native and mutant cellular αVβ3

To assess the contribution of the early contacts of 2.5D makes with αVβ3 on binding energy (Figure 5E), we replaced β3-Met^180^ with alanine (β3-Met^180^ has no homolog in β1) and β3-Arg^214^ with glycine (the equivalent residue in β_1_). MD simulations showed that the Met^180^/Arg^214^-Ala-Gly αVβ3 mutant (αVβ3^**^) sustained a significant loss in binding energy (25±19%) to 2.5D, but maintained the energy of interaction (111±19%) with 2.5F (Figure 6A, B). To validate the MD data, we quantified the binding of Alexa-647-labeled 2.5F and 2.5D to wild-type αVβ3 and αVβ3^**^, each transiently expressed in HEK293 cells. The double mutation reduced surface expression of αVβ3^**^ by ∼50% compared to wild-type αVβ3 (Figure 6C). When binding of each labeled ligand to the integrin was corrected for the degree of receptor expression, binding of 2.5F to αVβ3^**^ was minimally affected (85±9% of binding to wt-αVβ3, Figure 6D), but binding of 2.5D to αVβ3^**^ was markedly reduced (28±1% of binding to wt-αVβ3) (Figure 6E).

**Figure 6.**
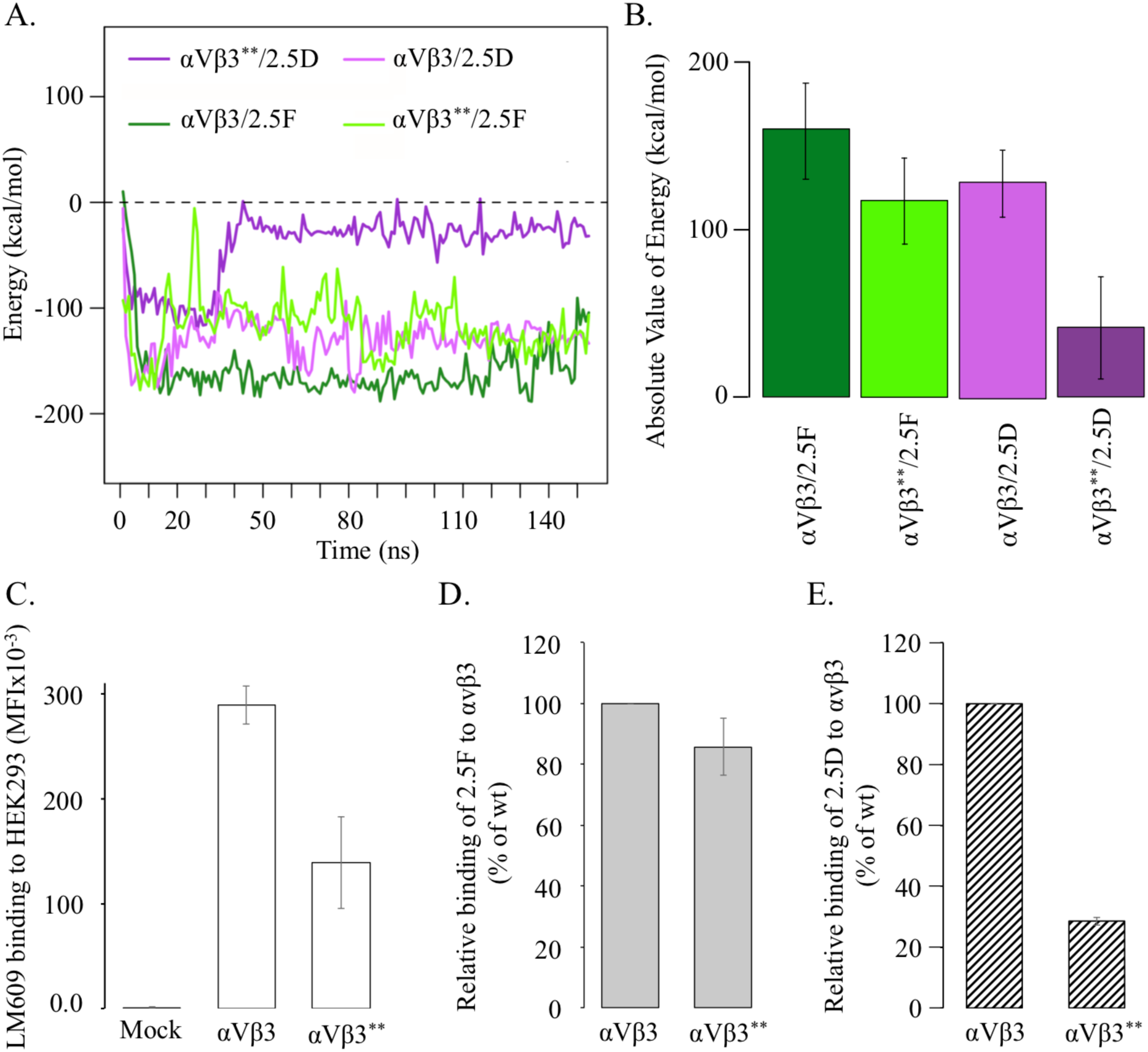
Interaction of 2.5D and 2.5F with wild type and mutant αVβ3. (A) MD simulations of 2.5D and 2.5F binding to wild type (WT) αVβ3 and αVβ3^**^ double mutant (β3-M180A and β3-R214G) showing energy over 160 ns. (B) Histograms showing values (mean± SD) of absolute energies from panel (A). Differences between αVβ3/2.5D and αVβ3**/2.5D and between αVβ3/2.5F and αVβ3**/2.5F were significant at p values of p<2.2×10^-16^ and 1.9×10^-6^, respectively. (C-E) Histograms (mean ± SD, n=4 independent experiments) showing binding of integrin antibody or Alexa 647-labelled 2.5F or 2.5D (each at 65 nM) to transiently transfected HEK293T-αVβ3 and double mutant β3-M180A and β3-R214G αVβ3^**^ in 1 mM Ca^2+^/Mg^2+^ as determined by FACS analysis. (C) Binding of αVβ3 heterodimer specific antibody LM609 detected by APC-labeled goat anti-mouse Fc-specific antibody. (D) Binding of Alexa647-2.5F to αVβ3 and αVβ3^**^. (E) Binding of Alexa647-2.5D to αVβ3 and αVβ3^**^. Binding of the knottin to wild-type αVβ3 in D and E for each experiment was set to 100%. No differences are seen in binding of 2.5F to WT or αVβ3^**^ (p>0.05) vs. 2.5D binding to the integrins (p=7.6×10^-8^).

## Discussion

The present studies show that specific recognition of αVβ3 by 2.5D requires high structural plasticity of the RGD-containing loop, revealed by comparing the conformational changes in loop backbone in structures of the apo-protein and αVβ3/2.5D complex. These comparisons also reveal a pronounced induced fit binding mechanism upon complex formation with αVβ3, which also resembles the well-known interactions between antibodies and antigens (Wilson and Stanfield, 1994). These features were not observed in binding of 2.5F to αVβ3, where the RGD-containing loop of the apo-protein is flexible, with some conformers having a 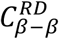 distance comparable to that found in the αVβ3-bound state, suggesting that 2.5F binds αVβ3 by conformation selection.

MD simulations elucidated the structural basis of the induced fit that underlies binding of 2.5D to αVβ3. The RGD-containing loop in the apo-protein is stabilized by a Type I β-turn, yielding a 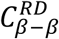 distance of ∼6.4Å, which extends to the optimal 9Å distance as a result of the switch of 2.5D-Trp^9^ from a solvent to a buried state. This switch appears to be driven by an early contacts with β3-Met^180^ and β3-Arg^214^, and is later influenced by the surrounding hydrophilic environment (β_3_-Tyr^166^, β3-Arg^214^), as 2.5D-Asp^8^ coordinates the metal ion at MIDAS. Substitution of β3-Met^180^ to Ala and β3-Arg^214^ to Gly (as in β_1_) resulted in a major loss in binding energy of 2.5D to αVβ3. This was confirmed by assessing knottin binding to HEK293 transiently expressing wild-type αVβ3 or αVβ3**. Since α5β1 lacks the equivalent Met and Arg residues, the induced fit mechanism cannot proceed, accounting for the lack of binding of 2.5D to α5β1. The 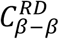 distance offered by some conformers of the flexible RGD-containing loop in 2.5F also accounts for its high affinity binding to α5β1.

The differences between 2.5F and 2.5D in adapting to the ligand-binding pocket in α5β1 likely relates to the RGD flanking residues of each ligand. Notably, 2.5F harbors two more prolines Pro^5^ and Pro^10^ in the RGD-containing loop (R<u>P</u>^5^*RGD*N<u>P</u>^10^P) when compared to that in 2.5D (Q<u>G</u>^5^*RGD*W<u>A</u>^10^P)(Figure 1A)(Kimura et al., 2009b). These trans-isomers of the prolines likely introduce local rigidity through their pyrrolidine ring, limiting the backbone dihedral angle to ∼ 90°, thus restraining the RGD-containing loop from adopting a β-turn fold, as in 2.5D (Krieger et al., 2005; Pabon and Camacho, 2017). Previous studies showed that the highly flexible linear RGD peptides have low affinity and are not specific for a particular integrin (Ruoslahti and Pierschbacher, 1987). Decreasing structural flexibility of the RGD loop by cyclization favors high affinity binding to integrins (Bogdanowich-Knipp et al., 1999), as it decreases the entropy term of the Gibbs free energy. However, our study shows that rigidifying the RGD loop limits the ability of 2.5D to bind certain integrins. The critical proline residues in the RGD-containing loop of 2.5F achieves a fine equilibrium between stability and flexibility of the RGD loop enabling a focused motional freedom (Krieger et al., 2005; Pabon and Camacho, 2017), where the RGD loop in 2.5F is flexible enough to bind both αVβ3 and α5β1 without excessive entropic contribution, which would hamper high affinity binding.

Eight of the 24 known mammalian integrins including αVβ3 and α5β1 bind to an RGD motif present in a host of natural ligands (Takada et al., 2007). High affinity peptidomimetics or RGD-like small molecules targeting single integrins have been developed. However, most of these continue to have residual but significant affinity to other RGD-binding integrins, and the selectivity of newer versions of these ligand-mimetics have not yet been fully explored in cell-based systems and at the high concentrations likely needed *in vivo* (Kapp et al., 2017). Given the ability of cancer cells to utilize both αVβ3 and α5β1 integrins for growth and metastasis, small molecules with multispecificity have been developed for potential applications in cancer therapy (Hatley et al., 2018; Sheldrake and Patterson, 2014). Despite these successes, development of multifunctional small molecule integrin antagonists that maintain high affinity and suitable pharmacokinetic properties remains a challenge (Nero et al., 2014). The engineered knottin miniproteins 2.5D and 2.5F have a number of advantages over small molecules and short peptides including exceptional structural stability, high affinity, and specificity to tumor-associated integrins (Kimura et al., 2009b; Kwan et al., 2017). Their amenability to large-scale synthesis provides a manufacturing advantage over monoclonal antibodies. These features highlight knottins as promising candidates to bridge the gap between small molecule drugs and monoclonal antibodies.

## Acknowledgements

This work was supported by NIH grants DK088327 and DK48549 (to MAA), DK101628 (to JVA), from the National Institutes of Diabetes, Digestive and Kidney diseases (NIDDK) of the National Institutes of Health; the Stanford Bio-X Interdisciplinary Initiatives Program, the Stanford Cancer Institute and the Stanford Maternal & Child Health Research Institute (JRC); National Science Foundation (NSF) grant CBET-0829205 (to MRKM). This work used resources of the Extreme Science and Engineering Discovery Environment (XSEDE) awarded to MRKM, which is supported by the National Science Foundation (NSF) (grant ACI-1053575). JRK was supported by a National Science Foundation Graduate Research Fellowship and an ARCS Scholar Award.

## Author contribution

JVA, JLA, JRK, BDA and JRC designed and performed the ligand-binding studies. FVC performed the NMR studies. JVA collected the x-ray diffraction data and refined the crystal structures with JPX. HS, KG and MRKM generated the molecular dynamic simulation data. All authors interpreted the data. MAA conceived and oversaw the project and wrote the manuscript with input from all authors.

## LEAD CONTACT AND MATERIALS AVAILABILITY

Further information and requests for resources and reagents should be directed to and will be fulfilled by the lead contact, M. Amin Arnaout.

## METHODS

### Peptide Synthesis

Knottins 2.5D, 2.5F and the scrambled FN-RDG2 (where RGD motif is replaced with RDG) were prepared as previously described (Kimura et al., 2009b). Briefly, the linear 33-amino acid peptides starting with Gly1 (Figure 1A) were made by solid-phase peptide synthesis on a CS Bio (Menlo Park, CA) instrument using standard 9-fluorenylmethyloxycarbonyl chemistry. Knottin peptides were folded by promoting disulfide bond formation in oxidizing buffer at room temperature with gentle rocking overnight. Folded knottins were purified by reversed-phase HPLC, where they appeared as a sharp peak with a shorter retention time than unfolded or misfolded precursors. The molecular masses of folded knottins were determined by matrix-assisted laser desorption/ionization time-of-flight (MALDI-TOF) mass spectrometry (Stanford Protein and Nucleic Acid Facility). Folded 2.5F and 2.5D (2 mg/mL) were incubated with an amine-reactive succinimidyl ester derivative of Alexa Fluor 647 carboxylic acid in 0.1 M Hepes, pH 8.0, at a 5:1 dye/peptide molar ratio for 1 h at room temperature and then at 4 °C overnight. The free dye was removed by dialysis and buffer exchange into phosphate buffered saline (PBS).

### Plasmids, mutagenesis, protein expression and purificatio

Human αVβ3 ectodomain was expressed in insect cells and purified as described (Mehta et al., 1998). The genetic sequence for knottins 2.5F, 2.5D or FN-RDG2 starting with Gly1 and ending with Gly33 (Figure 1A) was fused to the fragment crystallizable (Fc) region of mouse IgG2a in the pADD2 shuttle vector as described (Moore et al., 2013). The knottin Fc fusion proteins were expressed in human embryonic kidney (HEK293) cells following the manufacturer’s protocols in the FreeStyle MAX 293 Expression System (Invitrogen). Secreted knottin-Fc fusion proteins were purified using Protein A Sepharose (Sigma) followed by size exclusion chromatography (Superdex 75 column; GE Life Sciences). Purified knottin-Fc fusion proteins were bivalent homo-dimers of the expected molecular weight of **∼**60 kDa (Moore et al., 2013).

### Mammalian Cell lines

U87MG glioblastoma cells, K562 leukemia cells, and HEK293T embryonic kidney cells were obtained from American Type Culture Collection (Manassas, VA); integrin-transfected K562 cells (Blystone et al., 1994) were provided by Scott Blystone (SUNY Upstate Medical University).

### Binding assays

2.5D-Fc and 2.5F-Fc fusions were labeled with the succinimidyl ester derivative of Alexa Fluor 488 (Invitrogen) according to the manufacturer’s protocol. Free dye was removed by dialysis and buffer exchange into PBS. U87MG cells were detached using Enzyme-Free Cell Dissociation Buffer (Gibco); K562 cells were grown in suspension. 4×10^4^ cells then incubated with varying concentrations (0.01 – 500 nM) of Alexa Fluor 488 labeled 2.5D- or 2.5F-Fc fusion proteins in 25 mM Tris pH 7.4, 150 mM NaCl, 2 mM CaCl_2_, 1 mM MgCl_2_, 1 mM MnCl_2_, and 0.1% bovine serum albumin (BSA) for 3 hours at 4 °C, to minimize internalization. Cells were pelleted and washed twice with 800 µL of PBSA (phosphate buffered saline containing 0.1% bovine serum albumin) and the fluorescence of remaining surface-bound protein was measured using flow cytometry using a Guava EasyCyte 8HT instrument (EMD Millipore). Raw data were processed using FlowJo software (TreeStar Inc.).

Wild type αVβ3 and mutated αVβ3** transiently transfected HEK293T cells were gently trypsinized and washed in DPBS buffer. Cells were re-suspended in complete culture medium, incubated for one hour at 37°C and subsequently washed in 1 mM Ca^2+^/Mg^2+^, 0.1% bovine serum albumin-supplemented Hepes buffered saline pH 7.4 (binding buffer). 5×10^6^ cells were incubated with Alexa647-labeled 2.5F or 2.5D (50 nM) in binding buffer for 30 min, at 25 °C then washed, re-suspended, fixed in 1% paraformaldehyde and analyzed by flow cytometry in a LSRII flow cytometer (BD). Anti-α_V_β_3_ antibody LM609 (20 µg/ml) was used to normalize α_V_β_3_ cell surface expression in a separate set of tubes. Transfected HEK293T cells were stained with LM609 for 30 min at 4°C. After washing the excess antibody, APC-labeled goat anti-mouse Fc-specific antibody (10 µg/ml) was added for 30 min at 4°C, and the stained cells were washed, fixed and expression analyzed by flow cytometry as described above. Alexa-647 labeled knottin 2.5F or 2.5D binding to αVβ3 and αVβ3** was measured in mean fluorescence intensity (MFI) units, normalized according to LM609 binding and expressed as percentage of 2.5F or 2.5D binding to αVβ3.

### NMR

A codon-optimized DNA sequence was prepared by assembly PCR and cloned into the pET-32 vector to express a protein product in *E. coli* containing a thioredoxin and His-tag fusion protein separated by a TEV protease site. Uniform ^15^N- and ^13^C-labeling was achieved by IPTG-induced expression in BL21-DE3 cells in M9 minimal media containing ^15^NH_4_Cl and ^13^C-glucose. Cell lysis was followed by initial purification by His-tag capture with Ni-NTA. The thioredoxin fusion protein portion was removed with TEV protease to provide the exact 33-residue peptide sequence. Disulfide bond formation to fold the peptides was performed using the previously reported redox buffer (Kimura et al., 2009b). Final purification by RP-HPLC and characterization by ESI mass spectrometry confirmed folded engineered EETI-II peptides. NMR samples were prepared using sodium phosphate buffer, pH 6 containing 10% D_2_O. Standard multidimensional NMR datasets were acquired for backbone and side chain resonance assignments, along with ^13^C (aliphatic and aromatic) and ^15^N 3D-NOESY datasets for NOE-derived distance restraints. Dihedral angle restraints were derived from backbone assignments using TALOS+. 3D structure calculations were performed using the CYANA automated NOE assignment and simulated annealing algorithms. Initial structures calculated using standard automated NOE assignments led to convergence of 20 lowest energy structures, which were consistent with the expected knottin disulfide pattern. Further improved 3D structures were calculated by including disulfide bond restraints and hydrogen bond restraints determined by cross-hydrogen bond scalar couplings identified from long-range HNCO datasets. Final structures were refined by restrained molecular dynamics in explicit solvent using the YASARA package.

### Crystallography, structure determination and refinement

The αVβ3 ectodomain was crystallized at 4 °C by vapor diffusion using the hanging drop method as previously described (Xiong et al., 2009; Xiong et al., 2001; Xiong et al., 2002). Knottins 2.5F or 2.D (5 mM) was soaked into α_v_β_3_ crystals in the crystallization well solution containing 1 mM Mn^2+^ for 2–3 weeks. Crystals were harvested in 12% PEG 3500 (polyethylene glycol, molecular weight 3500) in 100 mM sodium acetate, pH 4.5, 800 mM NaCl plus 2 mM Mn^2+^ and 2.5F or 2.5D (at 5 mM), cryoprotected by the addition of glycerol in 2% increments up to a 24% final concentration and then flash frozen in liquid nitrogen. Diffraction data from cryo-cooled crystals were collected on the ID19 beamline fitted with a CCD detector at the APS Facility (Chicago, IL). Data were indexed, integrated and scaled with the HKL2000 (Otwinowski and Minor, 1997) program. Phases were determined by molecular replacement using PHASER (McCoy et al., 2007), with the structures α_V_β_3_ ectodomain (PDB ID 4MMX). Composite simulated annealing omit maps were then generated using the program Phenix Composite Omit Map package by turning on the simulated Cartesian annealing option with a temperature of 5,000K. The knottin structure was traced by the extra density using PDB 2IT7 and introducing the engineered mutations using Coot (Emsley and Cowtan, 2004). The resulting models were refined with the 1.10.1 version of Phenix (Adams et al., 2010) using simulated annealing, TLS, positional and individual temperature-factor refinement and default restrains. Several cycles of refinement and model building using Coot were applied to refine the structures of αVβ3/2.5D, αVβ3/2.5F (Table 2), with automatic optimization of X-ray and stereochemistry and additional Ramachandran restrains in the last cycles. A-weighted 2*fo-fc* electron density map was generated from the final models and structure factors using Phenix. All structural illustrations were prepared with the PyMol software (Schrödinger).

### Docking and Initial Configurations

The NMR structures of 2.5F (PDB: 6MM4) and 2.5D (PDB ID: 2M7T) were docked onto the crystal structures of the integrin αVβ3 (PDB ID: 4MMZ) and α5β1 (PDB ID: 4WJK) headpieces using the expert interface of the HADDOCK webserver. Four HADDOCK docking runs, between integrin α5β1 and 2.5D, integrin α5β1 and 2.5F, integrin αVβ3 and 2.5D, and integrin αVβ3 and 2.5F were performed. As docking inputs, the RGD sequence was specified as the active site of 2.5D and 2.5F. Residues α5-Glu^221^, α5-Asp^227^, and Mg^2+^ ion at the MIDAS site were specified as the active site of integrin α5β1, while residues αV-Asp^150^, αV-D^218^, and the MIDAS Mn^2+^ ion, were specified as the active site of integrin αVβ3. To preserve the ion coordination of the MIDAS, ADMIDAS, and LIMBS ions during the docking run, unambiguous distance restraints between the coordinating groups and the MIDAS, ADMIDAS, and LIMBS ions were fed into HADDOCK. Furthermore, integrins αVβ3 and α5β1 were specified as non-flexible in HADDOCK to prevent integrin backbone movement during docking. The RGD sequence of the knottins were specified as semi-flexible, while the non-RGD sequence was specified as non-flexible. Upon completion of the docking runs, the top solution from each generated cluster was analyzed. The best solution was then selected from these solutions based on maximal engagement of the specified active site residues.

### Molecular Dynamics Simulations

To simulate the interaction of 2.5D and 2.5F with integrins αVβ3 and α5β1, we performed Molecular Dynamics simulations using the top solution from each of the four HADDOCK structures. To setup the separated simulations, the axis between the center of mass of the integrin βA domain and the knottin variants was determined in the four HADDOCK complexes. Knottin was then separated from the integrin by 9Å along this axis, allowing water molecules to populate the space in between the knottin and the integrin upon solvation of the structure. We also performed equilibration simulations on the available crystal structures of αVβ3 in complex with 2.5D and 2.5F to be used as references. All structures were solvated and then ionized at a combined KCl concentration of 0.15 M. Structures were subsequently minimized for 100,000 steps and equilibrated for 0.5 ns using the NAMD molecular dynamics package (Phillips et al., 2005) and CHARMM27 force field (Brooks et al., 2009). Upon minimization, each of the four generated complexes ran for 500 ns. All simulations ran at an initial temperature of 310 K using the Nose-Hoover thermostat, and pressure was maintained at 1 atm using the Langevin piston. All equilibration and production run simulations were performed using a time step of 2 fs. Electrostatics of the system were determined using the Particle mesh Ewald (PME) method. van der Waals (VDW) interactions were modeled using a switching function to smoothly reduce the VDW force to zero at the cutoff distance of 1.2 nm. Simulations were then analyzed using Visual Molecular Dynamics (VMD) (Humphrey et al., 1996).

### Accession numbers

Coordinates for the NMR structures of knottins 2.5D and 2.5F and the crystal structures of αVβ3/2.5D and αVβ3/2.5F have been deposited in the PDB with ID codes 2M7T, 6MM4, 6MSL and 6MSU, respectively.

## EXPERIMENTAL MODEL AND SUBJECT DETAILS

Human glioblastoma cells (U87MG) were cultured in Dulbecco’s Modified Eagle Medium (DMEM) supplemented with 10% Fetal Bovine Serum (FBS) and 1% penicillin-streptomycin (P/S). K562 leukemia cells were grown in liquid culture in IMEM Iscove’s Modified Dulbecco’s Medium (IMDM) supplemented with 10% FBS and 1% P/S. Human embryonic kidney cells (HEK293T) were cultured in DMEM supplemented with 10% FCS, 2 mM l-glutamine, 1 mM sodium pyruvate, penicillin, and streptomycin, and were transiently co-transfected with pcDNA3 plasmids encoding full-length wild-type αVβ3, αVβ3** (αVβ3 with β3-M180A and β3-R214G) using Lipofectamine 2000 reagent (Invitrogen) according to the manufacturer’s protocol. 2.5D and 2.5F used for NMR structures were expressed in *E*.*coli* BL21 (DE3) cells grown in M9 minimal media containing ^15^NH_4_Cl and ^13^C-glucose.

## QUANTITATION AND STATISTICAL ANALYSIS

Equilibrium dissociation constants (*K*_*d*_) for the binding data shown I n Figures 1B and 1C were determined in SigmaPlot (Systat Software, San Jose, CA) using a least-square fit to a logistic curve. The standard error (SE) for *K*_*d*_ and t and p values were calculated using the reduced χ^2^ method in SigmaPlot. The validity of the *K*_*d*_ value was assessed with a p value cutoff of 0.05. In Fig. 2C, RMSD was calculated between the Cα’s in the different conformers and presented as. mean±SD in Fig. 2D. In Fig. 5B, the mean of the absolute values of the energies displayed in Fig. 5A are presented as histograms ± SE. p values were calculated with the Student’s t-test. In Fig. 6B, histograms derived from MD simulations of absolute energies shown in (A) are plotted ± SD, with p values calculated by the Student’s t-test. In Fig.6 C-E, values represent mean ±SD. p values were calculated with the Student’s t-test.

## DATA AND SOFTWARE AVAILABILITY

All software and libraries used are reported in the Method Details section, together with the Key Resources Table. Coordinates for the NMR structures of knottins 2.5D and 2.5F and the crystal structures of αVβ3/2.5D and αVβ3/2.5F have been deposited in the PDB with ID codes 2M7T, 6MM4, 6MSL and 6MSU, respectively.

