## Supplemental file for "Structural basis of the differential binding of engineered knottins to integrins αVβ3 and α5β1"

**Supplemental Information (SI)**

**Supplemental Figures (#3)**

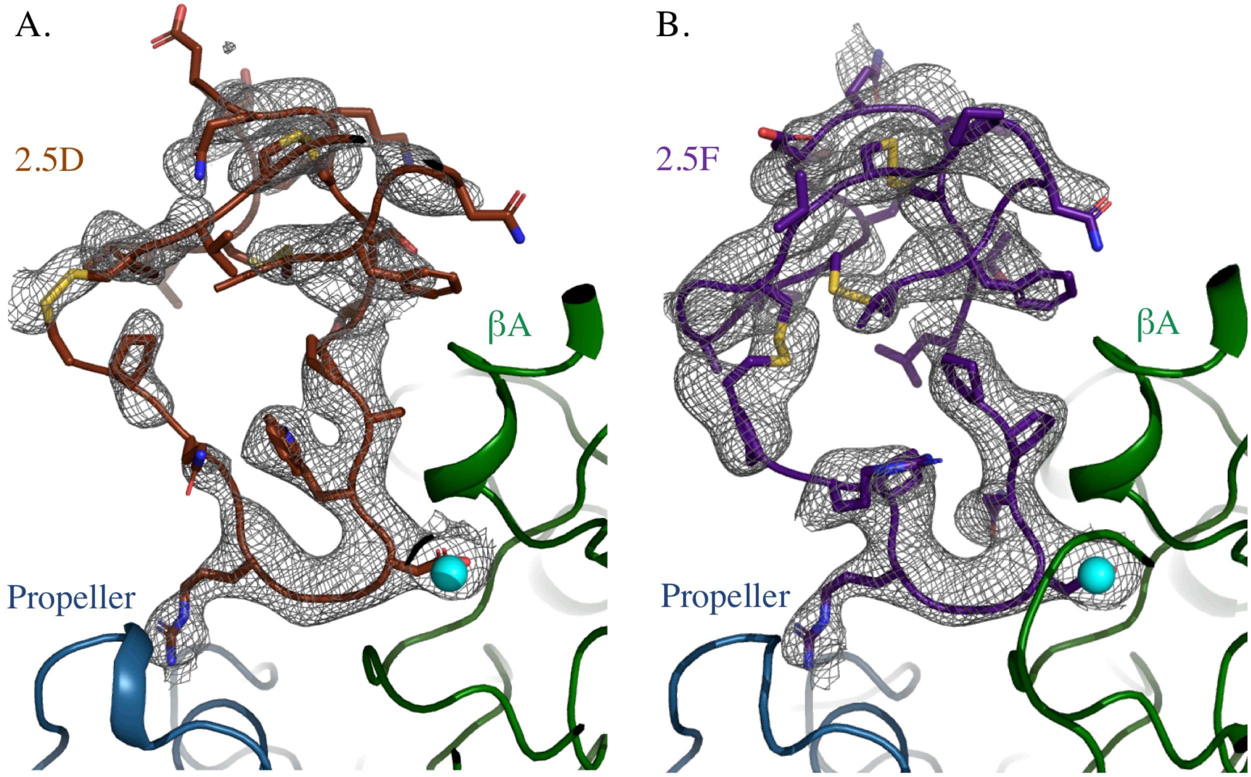


**Figure S1. Crystal structures of αVβ3 bound to 2.5D or 2.5F**. Composite simulated annealing omit maps (in grey isomesh) at 1.0 σ of 2.5D (in brown, A) and 2.5F (in (purple, B) in the knottin/αVβ3 structures (in ribbons) with only portions of the integrin propeller (blue) and βA (in green) domains shown. The MIDAS $Mn^{2+}$ ion is in cyan. Related to Figure 3.

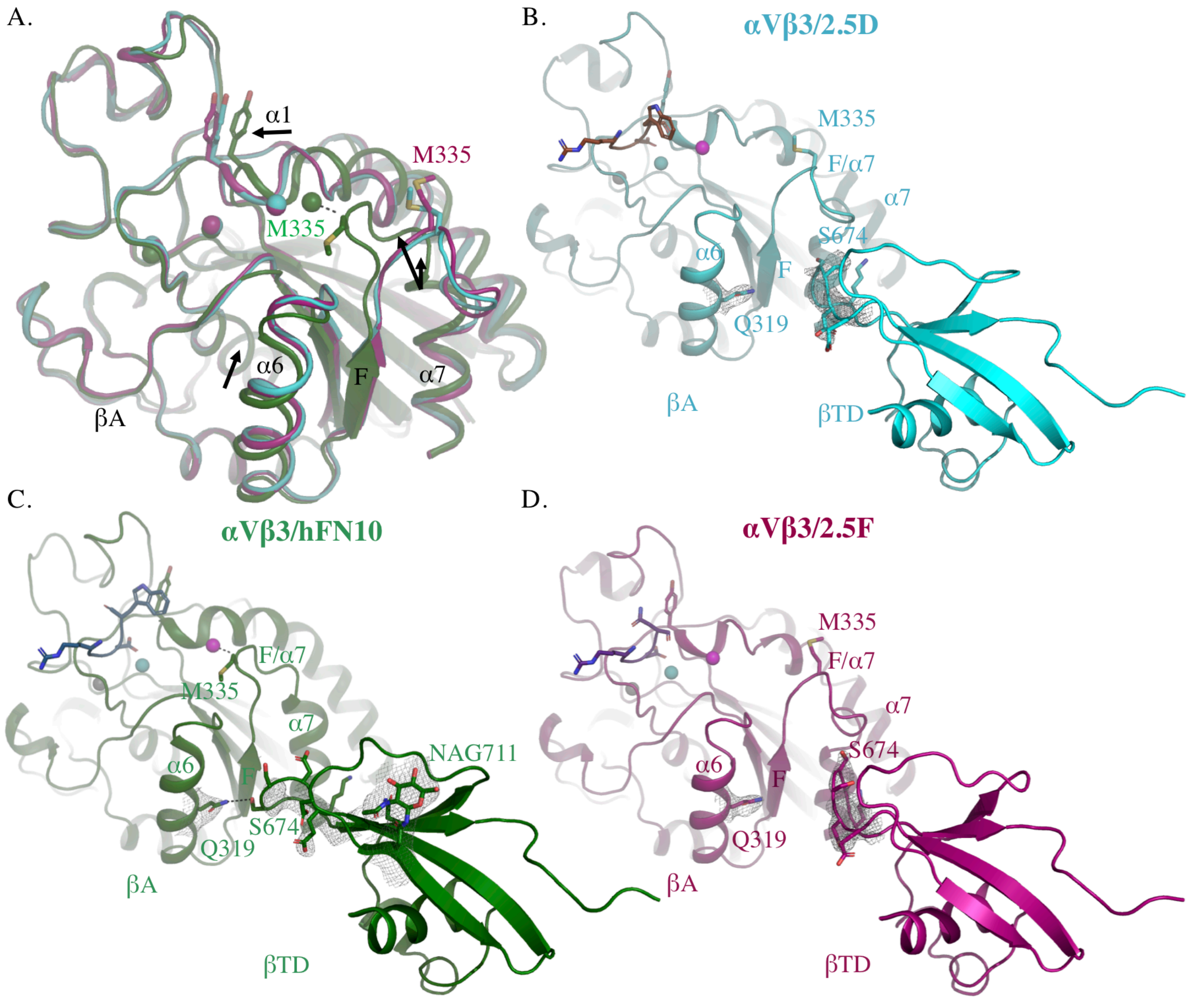


**Figure S2**: **Conformational changes and the βA/β-tail interface in crystal structures of αVβ3 in complex with 2.5D, 2.5F or hFN10.** A) Superimposition of βA domains of $\alpha_V\beta_3$ bound to hFN10 (green), 2.5F (dark pink) and 2.5D (cyan) shown in cartoon. Black arrows show the inward movement of βA-α1 towards MIDAS and translation of α6 between αVβ3/hFN10 (inactive) and αVβ3/2.5F or αVβ3/2.5D (active). LIMBS, MIDAS and ADMIDAS are shown in spheres. (B, C, D) Cartoon diagram of βA domain and β-tail domain (βTD) of αVβ3/2.5D (B), αVβ3/hFN10 (C), and αVβ3/2.5F (D). *2fo-fc* maps at 1.0 σ for βTD residues 671-676 and βA-Q319 in αVβ3/2.5D (B), αVβ3/hFN10 (C) and αVβ3/2.5F (D). NAG711 is shown in stick in αVβ3/hFN10 but is not detected in αVβ3/2.5D or αVβ3/2.5F. RGDW of 2.5D and hFN10, and RGDN of 2.5F are respectively shown in brown, light blue and purple sticks. LIMBS, MIDAS and ADMIDAS are shown in respectively grey, cyan and magenta spheres. Related to Figure 3.

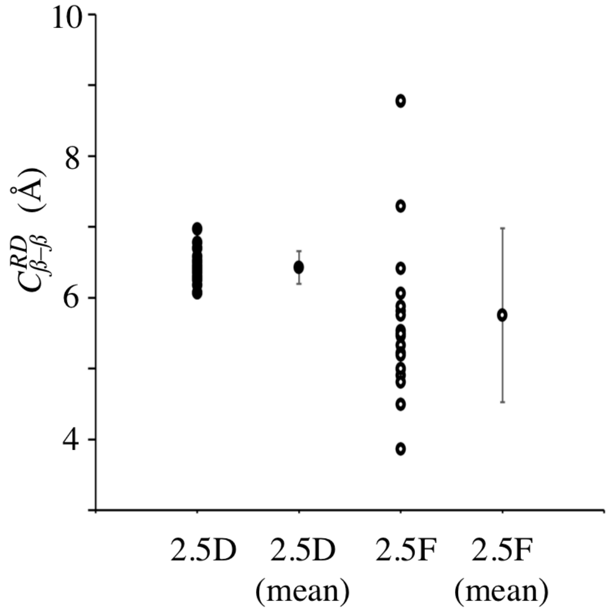


**Figure S3.** $\boldsymbol{C^{RD}_{\beta-\beta}}$ **in NMR structures of 2.5D and 2.5F.** Cβ-Cβ distance between $R^6$ and $D^8$ in each of the 20 NMR conformers of 2.5D (closed circles), and 2.5F (open circles). The respective mean values 6.42 ±0.23Å and 5.75 ±1.23Å are also shown. Related to Figure 4.

**Supplemental Movies (#4)**

**Supplemental Movie 1. 2.5F binding to αVβ3 in the trajectory from MD simulation in Fig 5C.** A ribbon representation of the backbone structure of 2.5F is colored in purple and portion of the αVβ3 head is colored in blue (propeller) and green (βA). Side chains for residues $R^6$ and $D^8$ of 2.5F, and $R^{214}$ and $M^{180}$ of β3 are shown in sticks with nitrogen in blue, oxygen in red and sulfur in yellow. Related to Figure 5.

**Supplemental Movie 2**. **2.5F binding to α5β1 in the trajectory from MD simulation in Fig 5D.** A ribbon representation of the backbone structure of 2.5F. Side chains for residues $R^6$ and $D^8$ and the integrin head are colored as in Movie 1. Related to Figure 5.

**Supplemental Movie 3**. **2.5D binding to αVβ3 in the trajectory from MD simulation in Fig 5E.** A ribbon representation of the backbone structure of 2.5D is colored in brown. Side chains of $R^6$, $D^8$ and $W^9$ of 2.5D, and $R^{214}$ and $M^{180}$ of βA are shown in sticks with nitrogen in blue, oxygen in red and sulfur in yellow. The integrin head is colored as in Movie 1. Related to Figure 5.

**Supplemental Movie 4. 2.5D binding to α5β1 in the trajectory from MD simulation in Fig 5F.** A ribbon representation of the backbone structure of 2.5D (in brown), and the portion of the α5β1 head colored as in Movie 1. Side chains for residues $R^6$, $D^8$ and $W^9$ of 2.5D are shown in sticks with nitrogen in blue, oxygen in red and sulfur in yellow. Related to Figure 5.
